# A fungal arrestin protein contributes to cell cycle progression and pathogenesis

**DOI:** 10.1101/801829

**Authors:** Calla L. Telzrow, Connie B. Nichols, Natalia Castro-Lopez, Floyd L. Wormley, J. Andrew Alspaugh

## Abstract

Arrestins, a structurally specialized and functionally diverse group of proteins, are central regulators of adaptive cellular responses in eukaryotes. Previous studies on fungal arrestins have demonstrated their capacity to modulate diverse cellular processes through their adaptor functions, facilitating the localization and function of other proteins. However, the mechanisms by which arrestin-regulated processes are involved in fungal virulence remain unexplored. We have identified a small family of four arrestins - Ali1, Ali2, Ali3, and Ali4 - in the human fungal pathogen *Cryptococcus neoformans*. Using complementary microscopy, proteomic, and reverse genetic techniques, we have defined a role for Ali1 as a novel contributor to cytokinesis, a fundamental cell cycle-associated process. We observed that Ali1 strongly interacts with proteins involved in lipid synthesis, and that *ali1*Δ mutant phenotypes are rescued by supplementation with lipid precursors that are used to build cellular membranes. From these data, we hypothesize that Ali1 contributes to cytokinesis by serving as an adaptor protein, facilitating the localization of enzymes that modify the plasma membrane during cell division, specifically the fatty acid synthases, Fas1 and Fas2. Finally, we assessed the contributions of the *C. neoformans* arrestin family to virulence, to better understand the mechanisms by which arrestin-regulated adaptive cellular responses influence fungal infection. We observed that the *C. neoformans* arrestin family contributes to virulence, and that the individual arrestin proteins likely fulfill distinct functions that are important for disease progression.

**IMPORTANCE:** To survive in unpredictable conditions, all organisms must adapt to stressors by regulating adaptive cellular responses. Arrestin proteins are conserved regulators of adaptive cellular responses in eukaryotes. Studies that have been limited to mammals and model fungi have demonstrated that disruption of arrestin-regulated pathways is detrimental for viability. The human fungal pathogen *Cryptococcus neoformans* causes more than 180,000 infection-related deaths annually, especially among immunocompromised patients. In addition to being genetically-tractable, *C. neoformans* has a small arrestin family of four members, lending itself to a comprehensive characterization of its arrestin family. This study serves as a functional analysis of arrestins in a pathogen, particularly in the context of fungal fitness and virulence. We investigate the functions of one arrestin protein, Ali1, and define its novel contributions to cytokinesis. We additionally explore the virulence contributions of the *C. neoformans* arrestin family and find that they contribute to disease establishment and progression.

## INTRODUCTION

Tight regulation of signal transduction pathways is necessary for appropriate cellular adaptation to the environment. Arrestins are a group of multifunctional proteins that modulate the activation and repression of diverse signaling pathways in eukaryotes (1–4). Structurally, arrestins are defined by protein domains with conserved β-sheet-rich regions, termed the N-terminal and C-terminal arrestin domains, that provide important secondary structure guiding protein localization and activity (5, 6). Functionally, arrestins link plasma membrane-initiated signals to intracellular responses by regulating signal internalization and intracellular signaling cascades. In doing so, arrestins enable the eukaryotic cell to fine-tune adaptive cellular responses through three specific mechanisms: desensitizing G protein-coupled receptors (GPCRs), scaffolding signaling cascades, and serving as adaptor proteins (1–4).

Nearly four decades ago, arrestins were first discovered for their unique ability to “arrest” cellular responses to persistent stimuli, in a classical process termed desensitization (7). Desensitization has been most commonly reported for and most extensively explored in visual and β-arrestins, classes of arrestins that are specific to metazoan cells (7–9). A third class of arrestins, the α-arrestins, are the evolutionary predecessors of the visual and β-arrestins (3, 5, 10–12). Present in all eukaryotes except for plants, α-arrestins share the ability to perform desensitization (13, 14). Beyond desensitization, non-traditional arrestin roles have been recently elucidated in model fungi. Fungal α-arrestins often act as scaffolds, physically bringing different components of signaling cascades within functional proximity of each other (10, 15–17). Additionally, other fungal α-arrestins function as adaptors, facilitating proper localization and function of other proteins. They often serve as ubiquitin ligase adaptors by means of proline-rich ubiquitin ligase binding motifs, or PxY sites, but α-arrestins can also act as adaptors for cytosolic proteins beyond those involved in ubiquitination (14, 18–21). Through these mechanisms, arrestins enable eukaryotic cells to terminate, promote, and modulate diverse adaptive cellular response signaling pathways both at the plasma membrane and throughout the cytosol.

The regulation of adaptive cellular responses is particularly important for pathogenic fungi because, in order to cause disease, they must quickly adjust to the hostile environment of the human host. Our laboratory and others have defined many fungal adaptive cellular response pathways, such as the Ras1 pathway and the Rim alkaline pH-sensing pathway, that are required for fungal virulence (22–27). However, the mechanisms by which adaptive cellular responses are regulated in pathogenic fungi are incompletely understood. The human fungal pathogen *Cryptococcus neoformans* is able to transition from its natural reservoir in the soil to establish infection in the host, resulting in more than 180,000 infection-related deaths annually, especially among immunocompromised patients (28). In contrast to other fungal model systems that encode numerous α-arrestin proteins in their genomes, we identified four α-arrestin proteins in *C. neoformans*: Ali1, Ali2, Ali3, and Ali4. This limited set of arrestin proteins allows for investigations of individual arrestin protein function, as well as assessment of arrestins as a collective family. Using Ali1 as a model, we report that Ali1 is a novel regulator of cytokinesis, and that this regulatory role is particularly important in the presence of stress. Additionally, we determine that Ali1 regulates cytokinesis through a typical arrestin role, likely functioning as an adaptor protein. Lastly, we demonstrate that, although Ali1 is not individually required for fatal infection, the α-arrestin family as a whole contributes to fungal virulence. By using the *C. neoformans* α-arrestins to explore the mechanisms by which fungal pathogens regulate their adaptive cellular responses, we can gain a deeper understanding of the establishment and progression of fungal infections.

## RESULTS

### *C. neoformans* contains a small family of four arrestin proteins

Previous work recently reported two putative α-arrestin proteins in the *C. neoformans* proteome, Ali1 (CNAG_02857) and Ali2 (CNAG_02341) (<u>A</u>rrestin-<u>Li</u>ke <u>1</u> and <u>2</u>) (Fig. 1) (25). These proteins were identified as α-arrestins based on the presence of the N-terminal and C-terminal arrestin domains. We performed a search of the *C. neoformans* proteome to identify all α-arrestin domain-containing proteins (29). In doing so, we identified two additional α-arrestin proteins, Ali3 (CNAG_ 04137) and Ali4 (CNAG_ 05343), each of which contains a single C-terminal arrestin domain (Fig. 1). In addition to the arrestin domains, each of the identified *C. neoformans* α-arrestin proteins also contains multiple ubiquitin ligase binding sites, or PxY sites, which are common features of α-arrestins (Fig. 1) (3, 12, 14). For the sake of simplicity, the *C. neoformans* α-arrestins will simply be referred to as “arrestins” throughout the remainder of this manuscript.

**FIGURE 1.**
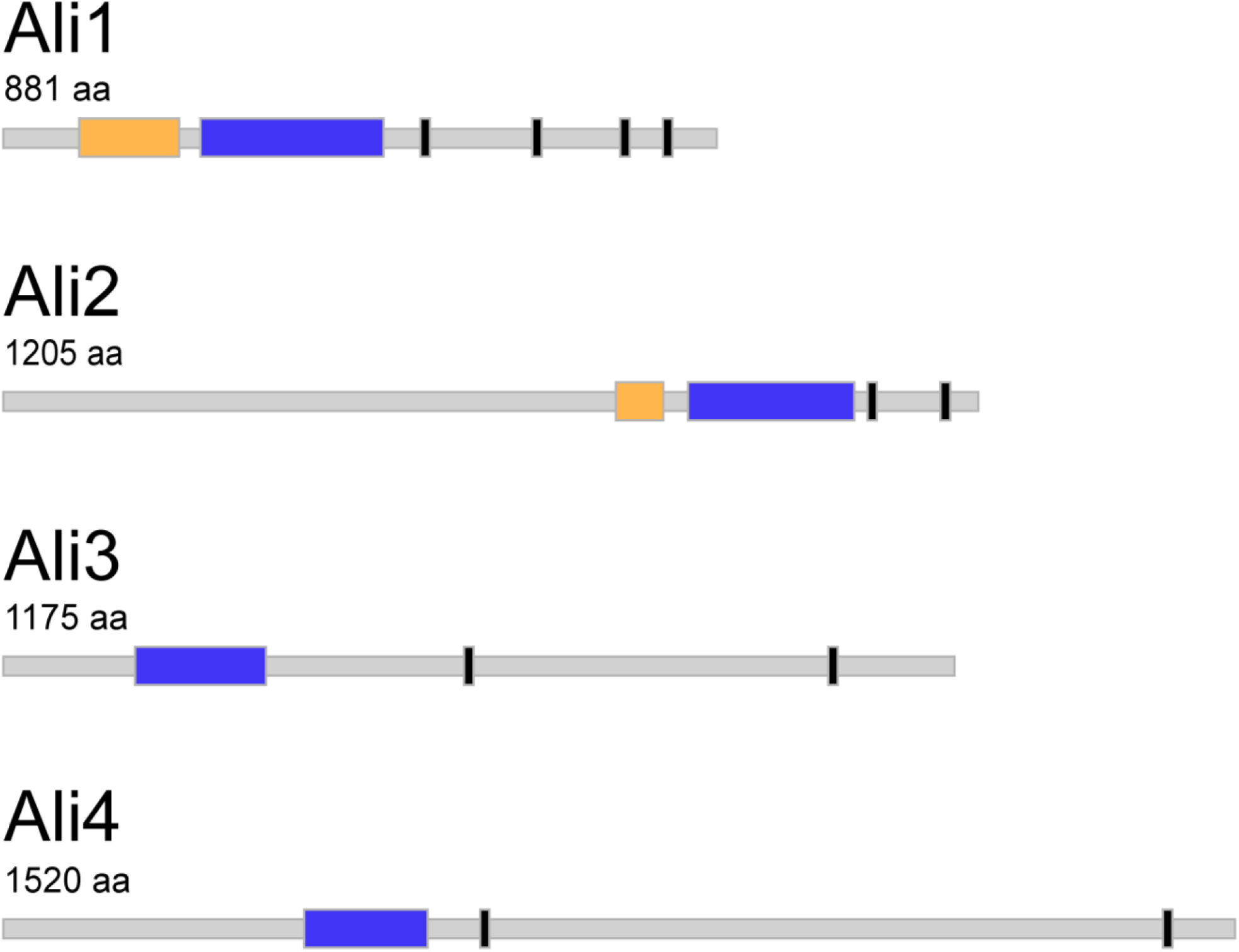
The *C. neoformans* arrestin proteins. The arrestin proteins within the *C. neoformans* proteome – Ali1, Ali2, Ali3, and Ali4 – were identified by the presence of the conserved β-sheet-rich arrestin domains. If present, the N-terminal arrestin domain (yellow), the C-terminal arrestin domain (blue), and any potential ubiquitin ligase binding sequences, or PxY sequences (black), are indicated for each arrestin protein. Protein and domain sizes are depicted to scale (aa = amino acids).

To prioritize our studies, we compared the protein sequence of each of the *C. neoformans* arrestin proteins with those in *Saccharomyces cerevisiae* and humans, two organisms with well-characterized arrestin families. Because there is often limited protein sequence conservation between arrestins in different species, we used two different programs within the Basic Local Alignment Search Tool (BLAST) algorithm. Protein-protein BLAST (blastp) was utilized to detect arrestin proteins in the *S. cerevisiae* and human proteomes with moderate to high degrees of homology with the *C*. *neoformans* arrestin proteins (30). We also used Position-Specific Iterated BLAST (PSI-BLAST) to detect arrestin proteins in the *S. cerevisiae* and human proteomes with low, but potentially relevant, degrees of homology with the *C. neoformans* arrestin proteins (31, 32). Ali1 was the only *C. neoformans* arrestin protein that shared significant sequence homology with multiple *S. cerevisiae* arrestins and a human arrestin (Tables S1 & S2). In all of these instances, the identified sequence homology was located within the arrestin domains of both proteins. We therefore elected to focus our initial studies on Ali1.

### Ali1 exhibits cell cycle-regulated localization that is dependent on the Ras signaling pathway

We first investigated the subcellular localization of Ali1, positing that its localization would be indicative of function. We C-terminally tagged Ali1 with green fluorescent protein (GFP) and validated proper expression, stability, and function of the Ali1-GFP fusion protein using quantitative real time PCR, western blotting, and mutant phenotype complementation, respectively (data not shown). Following validation, we incubated the wild-type (WT) strain and the Ali1-GFP strain to mid-logarithmic growth phase in yeast-peptone-dextrose (YPD) medium at 30°C, a nutrient-rich growth condition, and tissue culture (TC) medium at 37°C, a stressful condition that more closely mimics the host environment. Using epifluorescence microscopy, we observed identical patterns of localization in both conditions: Ali1-GFP localizes diffusely throughout the cytoplasm and is excluded from the vacuole in non-budding cells (Fig. 2A). However, in budding cells, Ali1-GFP is enriched at the developing septum, and it also localizes within discrete puncta at the poles (Fig. 2A). To confirm this localization, we performed subcellular fractionations to measure the relative abundances of Ali1-GFP within the soluble (cytoplasmic) and insoluble (membrane-associated) cellular fractions. We observed that Ali1-GFP is enriched in the insoluble fraction, indicating that Ali1-GFP is associated with insoluble cellular components such as the plasma membrane, intracellular membranes, and cell wall components (Fig. 2B). Together these observations suggest that Ali1 may be a novel contributor to cell polarity and/or cell division.

**FIGURE 2.**
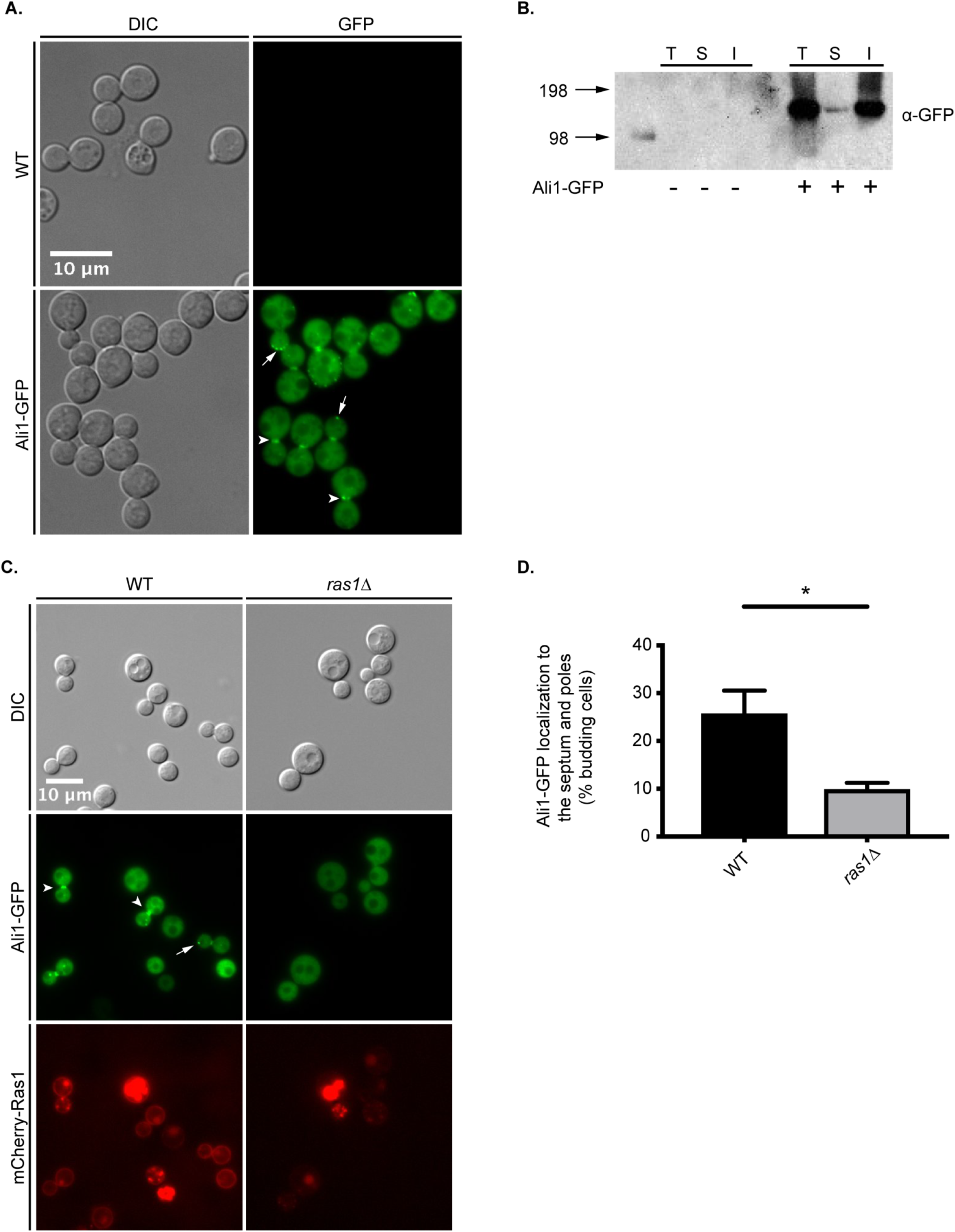
Ali1 subcellular localization patterns. A. The WT and Ali1-GFP strains were incubated in YPD medium at 30°C (above) or TC medium at 37°C, and Ali1-GFP was localized by epifluorescence microscopy (Zeiss Axio Imager A1). Ali1-GFP localization to the septum (arrowheads) and poles (arrows) of budding yeasts is depicted. B. To determine the relative enrichment of Ali1-GFP in different cellular fractions, WT and Ali1-GFP total cell lysates (T) were subjected to ultracentrifugation (30,000 × g) to isolate the soluble (S) and insoluble (I) cellular fractions. Samples were analyzed by western blotting using an anti-GFP antibody. The estimated size of Ali1-GFP is approximately 122 kDa. C. The dependence of Ali1-GFP localization on the Ras1 signaling pathway was determined using galactose-inducible expression of the *RAS1* transcript. Cells were incubated in YPGal (WT) or YPD (*ras1*Δ) media. Ali1-GFP localization to the septum (arrowheads) and poles (arrows) of budding yeasts was observed using epifluorescence microscopy (Zeiss Axio Imager A1). D. The frequency of Ali1-GFP localization to the septum and poles was quantified in the presence and absence of Ras1. The percentage of actively budding cells that displayed Ali1-GFP localization to the septum and/or poles was calculated in both YPGal (WT) and YPD (*ras1*Δ) conditions. A minimum of 600 cells were analyzed in both YPGal (WT) and YPD (*ras1*Δ) conditions across three biological replicates (n = 3). Error bars represent the standard error of the mean (SEM). Log transformation was used to normally distribute the data for statistical analysis (*Student’s *t*-test p < 0.05).

Previous work in our group identified *C. neoformans* Ras1 as a GTPase that is required for cytokinesis and polarized growth, particularly in the presence of cell stress (22, 23, 33). Therefore, we hypothesized that Ras1 might also be required for the cell cycle-associated localization of Ali1-GFP. To test this, we constructed a strain that, in addition to expressing *ALI1-GFP*, also expressed *mCherry-RAS1* under a galactose-regulatable promoter (23). When incubated in galactose as the sole carbon source (YPGal), cells express *RAS1* at levels similar to WT cells, which is confirmed by mCherry-Ras1 localization to the plasma membrane. In contrast, *RAS1* expression is repressed when this strain is incubated in glucose as the sole carbon source (YPD). When incubated in YPGal (WT) conditions, Ali1-GFP localizes to the septum and poles of dividing cells as previously observed (Fig. 2C). However, in YPD (*ras1*Δ) conditions, Ali1-GFP localization to the septum and poles of budding cells is impaired (Fig. 2C). We quantified the frequency of Ali1-GFP localization to the septum and poles specifically among budding cells and observed that the polarized pattern of Ali1-GFP localization is significantly decreased in YPD (*ras1*Δ) conditions compared to YPGal (WT) conditions (Fig. 2D). These data indicate that Ali1-GFP localization to sites associated with cell polarity is dependent on Ras1.

### Ali1 is a regulator of cytokinesis

From the distinct, cell cycle-regulated localization pattern of Ali1-GFP, we hypothesized that Ali1 is involved in the process of cell division. To test this hypothesis, we constructed a loss-of-function *ali1*Δ mutant and analyzed this strain for cytokinesis defects. *C. neoformans* cells with loss-of-function mutations in septin genes, known contributors to cytokinesis, exhibit cytokinesis defects when grown at elevated temperatures (34). We incubated the WT strain, the *ali1*Δ mutant, and the complemented (*ali1*Δ + *ALI1*) strain at the permissive temperature of 30°C, or the more stressful temperature of 39°C, and assessed the cells for cytokinesis defects by DIC microscopy. We observed that the *ali1*Δ mutant exhibits similar morphology to the WT strain at 30°C (Fig. 3A). However, at 39°C, we observed that the *ali1*Δ mutant displays an increased incidence of cytokinesis defects, specifically elongated cells, wide bud necks, and cells that fail to complete cytokinesis (Fig. 3A) (23, 34). We quantified this observation and found that the *ali1*Δ mutant exhibits a higher frequency of cytokinesis defects at 39°C than the WT strain, and that this *ali1*Δ mutant phenotype is rescued by complementation with the WT *ALI1* allele (Fig. 3B & S1). This observation implicates Ali1 in the regulation of cytokinesis.

**FIGURE 3.**
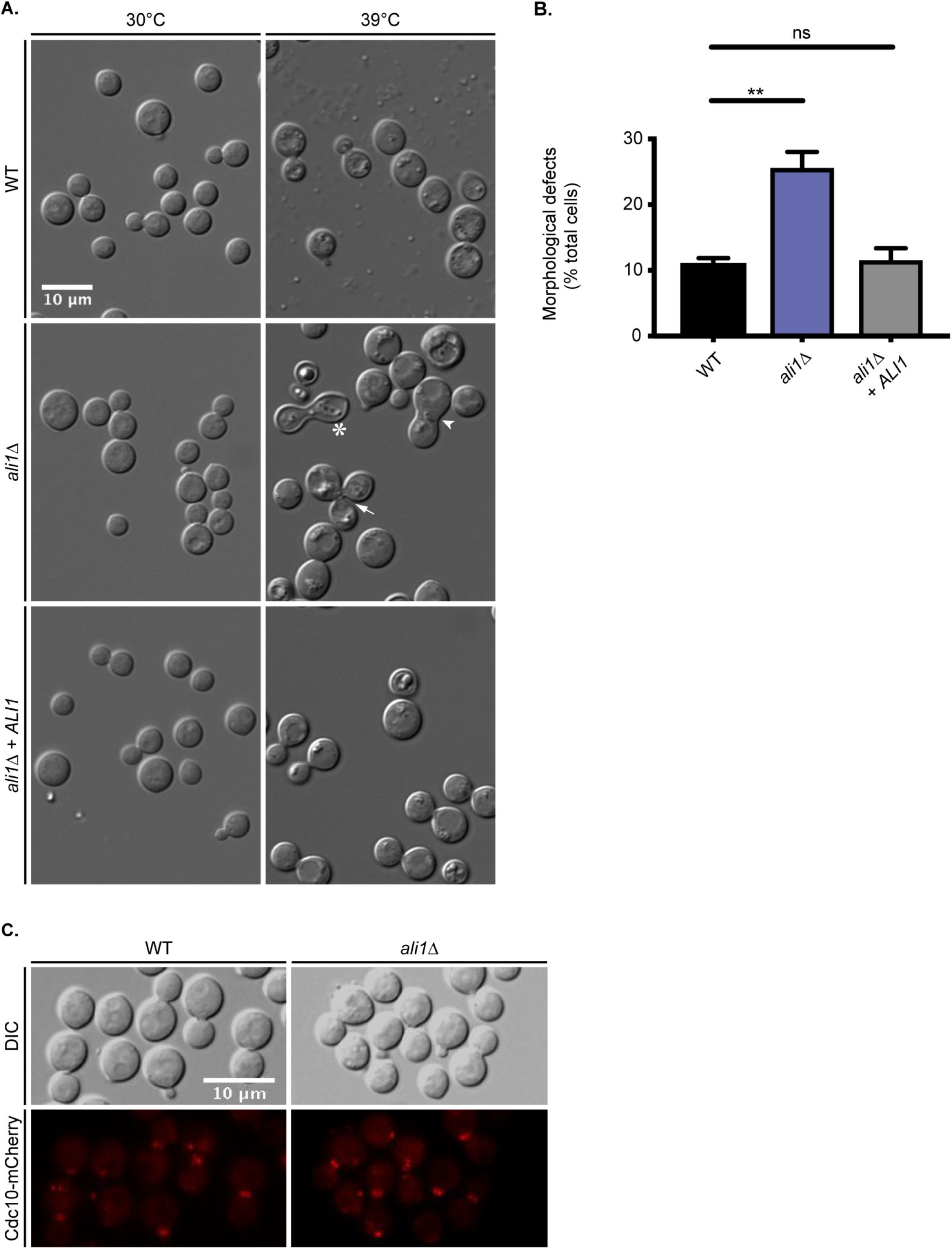
Cellular morphology of the *ali1*Δ mutant. A. The WT, *ali1*Δ mutant, and *ali1*Δ + *ALI1* strains were incubated in YPD medium at either 30°C or 39°C and subsequently imaged by DIC microscopy (Zeiss Axio Imager A1). The *ali1*Δ mutant cells displaying morphological defects, such as elongated cells (asterisk), wide bud necks (arrowhead), and cytokinesis failure (arrow), are indicated. B. The percentage of total cells displaying morphological defects at 39°C was quantified for each strain. A minimum of 600 cells were analyzed across three biological replicates (n = 3). Error bars represent the SEM. Log transformation was used to normally distribute the data for statistical analysis (**One-way ANOVA p < 0.01; ns = not significant). C. The septin protein, Cdc10, was localized by visualization of the Cdc10-mCherry fusion protein in both the WT and the *ali1*Δ mutant backgrounds after incubation in YPD medium at 30°C (above) or 37°C. The Cdc10-mCherry fusion protein was localized using epifluorescence microscopy (Zeiss Axio Imager A1).

Because the temperature-dependent cytokinesis phenotype of the *ali1*Δ mutant mimics that of septin mutants, we next hypothesized that Ali1 is required for septin protein complex formation. To do so, we analyzed septin protein localization in the WT strain compared to the *ali1*Δ mutant. Using epifluorescence microscopy, we observed that the septin protein, Cdc10-mCherry, localizes to the septum of budding cells in both the WT and *ali1*Δ mutant backgrounds, indicating that Ali1 is not required for assembly of septin proteins at the site of septum formation (Fig. 3C). Although Ali1 is not required for this particular septin protein localization, we next hypothesized that Ali1 may be involved in cytokinesis by modulating the function of other interacting proteins at the septum and poles.

### Ali1-GFP interacts with proteins involved in lipid metabolism

To better understand the mechanism by which Ali1 regulates cytokinesis, we performed a proteomic screen to identify potential protein interactors of Ali1. To do so, we incubated the Ali1-GFP strain, and the WT strain as a negative control, to mid-logarithmic growth phase in YPD medium. We subsequently conditioned the cultures in YPD or TC media at 30°C, in order to capture protein-protein interactions at the most permissive temperature, for three hours. Following cell lysis, GFP immunoprecipitations were performed to enrich for Ali1-GFP. The immunoprecipitations were then analyzed by LC/ESI/MS/MS to identify proteins that potentially interact with Ali1-GFP in these two conditions. A total of 1,122 proteins were identified as potential Ali1-GFP interactors using this approach (Table S3). We applied unbiased methods to enrich for proteins in both YPD and TC conditions that were highly represented in the Ali1-GFP immunoprecipitations and lowly represented, if at all, in the respective WT immunoprecipitation. This prioritization scheme resulted in 59 and 62 potentially biologically-relevant protein interactors of Ali1-GFP in YPD and TC conditions, respectively (Tables S4 & S5). Table 1 displays the top 30 hits from this experiment in YPD medium, organized by decreasing average exclusive unique peptide count (APC) and increasing APC indentified in the WT immunoprecipitation. Table 2 displays the top 30 hits from this experiment in TC medium, organized by decreasing APC and increasing APC indentified in the WT immunoprecipitation.

**TABLE 1.** The most abundant 30 proteins identified as potential Ali1-GFP interactors in YPD medium*^a^*

| Gene ID | Gene Name | Gene Product | Average Peptide Count (APC) | Percent of APC identified in WT |
| --- | --- | --- | --- | --- |
| CNAG_02099 | *FAS1* | Fatty acid synthase, beta subunit | 9.0 | 0.0 |
| CNAG_02100 | *FAS2* | Fatty acid synthase, alpha subunit | 8.7 | 11.5 |
| CNAG_02748 |  | UTP-glucose-1-phosphate uridylyltransferase | 8.7 | 11.5 |
| CNAG_07373 |  | Carbamoyl-phosphate synthase, large subunit | 8.0 | 0.0 |
| CNAG_03944 |  | Chaperone regulator | 6.7 | 15.0 |
| CNAG_03358 |  | Phosphoglycerate kinase | 6.0 | 0.0 |
| CNAG_04441 |  | Polyadenylate-binding protein, cytoplasmic and nuclear | 6.0 | 16.7 |
| CNAG_01464 | *FHB1* | Flavohemoglobin | 5.7 | 17.6 |
| CNAG_00418 |  | S-adenosylmethionine synthase | 5.3 | 18.8 |
| CNAG_01586 |  | F-type H-transporting ATPase, B subunit | 5.0 | 0.0 |
| CNAG_02545 |  | Inorganic pyrophosphatase | 5.0 | 0.0 |
| CNAG_04659 |  | Pyruvate decarboxylase | 4.7 | 0.0 |
| CNAG_02928 |  | Large subunit ribosomal protein L5e | 4.3 | 0.0 |
| CNAG_05759 |  | Acetyl-CoA carboxylase/biotin carboxylase | 4.0 | 0.0 |
| CNAG_07363 |  | Isocitrate dehydrogenase, NAD-dependent | 4.0 | 0.0 |
| CNAG_02673 |  | NAD dependent epimerase/dehydratase | 3.7 | 0.0 |
| CNAG_02811 |  | Small subunit ribosomal protein S29 | 3.7 | 0.0 |
| CNAG_07745 | *MPD1* | Alcohol dehydrogenase, propanol-preferring | 3.3 | 0.0 |
| CNAG_00176 |  | Glutamate carboxypeptidase | 3.3 | 0.0 |
| CNAG_01404 |  | Hsp71-like protein | 3.3 | 0.0 |
| CNAG_02500 |  | Calnexin | 3.0 | 0.0 |
| CNAG_02991 |  | Cofilin | 3.0 | 0.0 |
| CNAG_02943 |  | Cytoplasmic protein | 3.0 | 0.0 |
| CNAG_03588 | *LYS2* | L-aminoadipate-semialdehyde dehydrogenase | 3.0 | 0.0 |
| CNAG_02736 |  | T-complex protein 1, theta subunit | 3.0 | 0.0 |
| CNAG_03765 | *TPS2* | Trehalose-phosphatase | 3.0 | 0.0 |
| CNAG_00136 |  | Ubiquitin-activating enzyme E1 | 3.0 | 0.0 |
| CNAG_07558 |  | Uncharacterized protein | 3.0 | 0.0 |
| CNAG_06112 |  | Carbamoyl-phosphate synthase arginine-specific, large chain | 2.7 | 0.0 |
| CNAG_00879 |  | Glutamate dehydrogenase | 2.7 | 0.0 |

<sup>a</sup> The average peptide count (APC) was calculated by averaging the exclusive unique peptide counts for each potential interactor across three biological replicates. The percent of APC identified in the WT immunoprecipitation was calculated by dividing the

**TABLE 2.** The most abundant 30 proteins identified as potential Ali1-GFP interactors in TC medium*^a^*

| Gene ID | Gene Name | Gene Product | Average Peptide Count (APC) | Percent of APC identified in WT |
| --- | --- | --- | --- | --- |
| CNAG_02099 | *FAS1* | Fatty acid synthase, beta subunit | 19.3 | 0.0 |
| CNAG_02100 | *FAS2* | Fatty acid synthase, alpha subunit | 16.7 | 6.0 |
| CNAG_04327 |  | Uncharacterized protein | 10.3 | 9.7 |
| CNAG_05355 | *RSP5* | E3 ubiquitin-protein ligase | 6.0 | 0.0 |
| CNAG_05978 |  | Glutamate-tRNA ligase | 6.0 | 16.7 |
| CNAG_03588 | *LYS2* | L-aminoadipate-semialdehyde dehydrogenase | 6.0 | 16.7 |
| CNAG_03701 |  | 3-phosphoshikimate 1-carboxyvinyltransferase | 5.7 | 17.6 |
| CNAG_00743 |  | Imidazoleglycerol phosphate synthase, cyclase subunit | 5.3 | 0.0 |
| CNAG_07561 |  | 6-phosphogluconate dehydrogenase, decarboxylating I | 5.3 | 18.8 |
| CNAG_06175 |  | 26S proteasome, regulatory subunit N2 | 4.7 | 0.0 |
| CNAG_01216 |  | 6-phosphogluconolactonase | 4.0 | 0.0 |
| CNAG_00602 |  | Eukaryotic translation initiation factor 3, subunit I | 4.0 | 0.0 |
| CNAG_00136 |  | Ubiquitin-activating enzyme E1 | 4.0 | 0.0 |
| CNAG_06666 |  | Alpha-1,4 glucan phosphorylase | 3.7 | 0.0 |
| CNAG_02565 |  | Homoaconitase, mitochondrial | 3.7 | 0.0 |
| CNAG_02035 |  | Triosephosphate isomerase | 3.7 | 0.0 |
| CNAG_05650 | *UBP5* | Ubiquitin carboxyl-terminal hydrolase 7 | 3.7 | 0.0 |
| CNAG_00708 |  | Pre-mRNA-splicing factor Slt11 | 3.3 | 0.0 |
| CNAG_01981 |  | Sulfide:quinone oxidoreductase | 3.3 | 0.0 |
| CNAG_03249 |  | THO complex, subunit 4 | 3.3 | 0.0 |
| CNAG_00649 |  | Tryptophan synthase, beta subunit | 3.3 | 0.0 |
| CNAG_02858 | *ADE12* | Adenylosuccinate synthetase | 3.0 | 0.0 |
| CNAG_01189 |  | DNA-directed RNA polymerase I, subunit RPA1 | 3.0 | 0.0 |
| CNAG_02545 |  | Inorganic pyrophosphatase | 3.0 | 0.0 |
| CNAG_04976 |  | Zuotin | 3.0 | 0.0 |
| CNAG_04951 |  | 3-deoxy-7-phosphoheptulonate synthase | 2.7 | 0.0 |
| CNAG_02500 |  | Calnexin | 2.7 | 0.0 |
| CNAG_06730 | *GSK3* | CMGC/GSK protein kinase | 2.7 | 0.0 |
| CNAG_00700 |  | Phosphoribosylaminoimidazolecarboxamide formyltransferase/IMP cyclohydrolase | 2.7 | 0.0 |
| CNAG_02315 |  | Ubiquinol-cytochrome c reductase, iron-sulfur subunit | 2.7 | 0.0 |

<sup>a</sup> The average peptide count (APC) was calculated by averaging the exclusive unique peptide counts for each potential interactor across three biological replicates. The percent of APC identified in the WT immunoprecipitation was calculated by dividing the

We observed that the Ali1-GFP protein interactome, both in YPD and TC conditions, was enriched in proteins involved in two biological processes: protein localization/stability and lipid metabolism. Enrichment of proteins involved in protein localization/stability, such as ubiquitination proteins, has been reported for arrestins in other organisms, particularly arrestins that perform adaptor functions (1, 11, 12, 35). Ubiquitination proteins were found in both conditions, but were more highly represented in TC conditions along with various proteasome subunits (Tables 1 & 2; Tables S4 & S5). Supporting our Ali1-GFP localization observations, the septin proteins Cdc10, Cdc11, and Cdc12 were also identified at low abundances, indicating potential transient interactions with Ali1-GFP (Table S3). Interestingly, in addition to protein localization/stability, the interactome of Ali1-GFP was highly enriched in proteins involved in lipid metabolism. Multiple proteins involved in lipid synthesis and degradation were identified in both YPD and TC conditions. Specifically, the fatty acid synthase β subunit, Fas1, was the overall strongest potential interactor in both conditions (Tables 1 & 2). The enzymatic partner of Fas1, the fatty acid synthase α subunit, Fas2, was also identified at very high abundances in both YPD and TC conditions (Tables 1 & 2). The observation that Fas1 and Fas2 were the most abundant interactors in multiple, independent experiments conducted in both YPD and TC conditions, as well as the fact that previous proteomic experiments we have conducted with other proteins of interest did not find enrichment of the fatty acid synthases, suggest that Fas1 and Fas2 are true, specific interactors of Ali1 (26). These data indicate that Ali1 may be involved in the regulation of localized lipid production at the developing septum and poles of budding cells, assisting in efficient cytokinesis, especially in stressful growth conditions.

### The *ali1*Δ mutant has impaired cell surface integrity that is rescued by lipid precursor supplementation

Using the data collected from the Ali1-GFP proteomic screen, we further explored the mechanism by which Ali1 regulates cytokinesis. As previously discussed, we identified many potential interactors of Ali1-GFP involved in lipid metabolism, a process that is essential for proper synthesis and organization of the cell surface, specifically the cell membrane. Therefore, we assessed the cell surface integrity of the *ali1*Δ mutant. We incubated the WT strain, the *ali1*Δ mutant, and the *ali1*Δ + *ALI1* strain at 30°C in the presence of various cell surface stressors: calcofluor white, Congo red, SDS, and caffeine (36–39). The *ali1*Δ mutant exhibits modest susceptibility to caffeine, a cell surface stressor that serves as a marker of cell surface integrity, when incubated at 30°C (Fig. 4A) (40, 41). This phenotype is drastically enhanced when the *ali1*Δ mutant is incubated at the more stressful temperature of 37°C (Fig. 4B). At both temperatures, this sensitivity is rescued by complementation with the WT *ALI1* allele, indicating likely alterations to the *ali1*Δ mutant cell surface.

**FIGURE 4.**
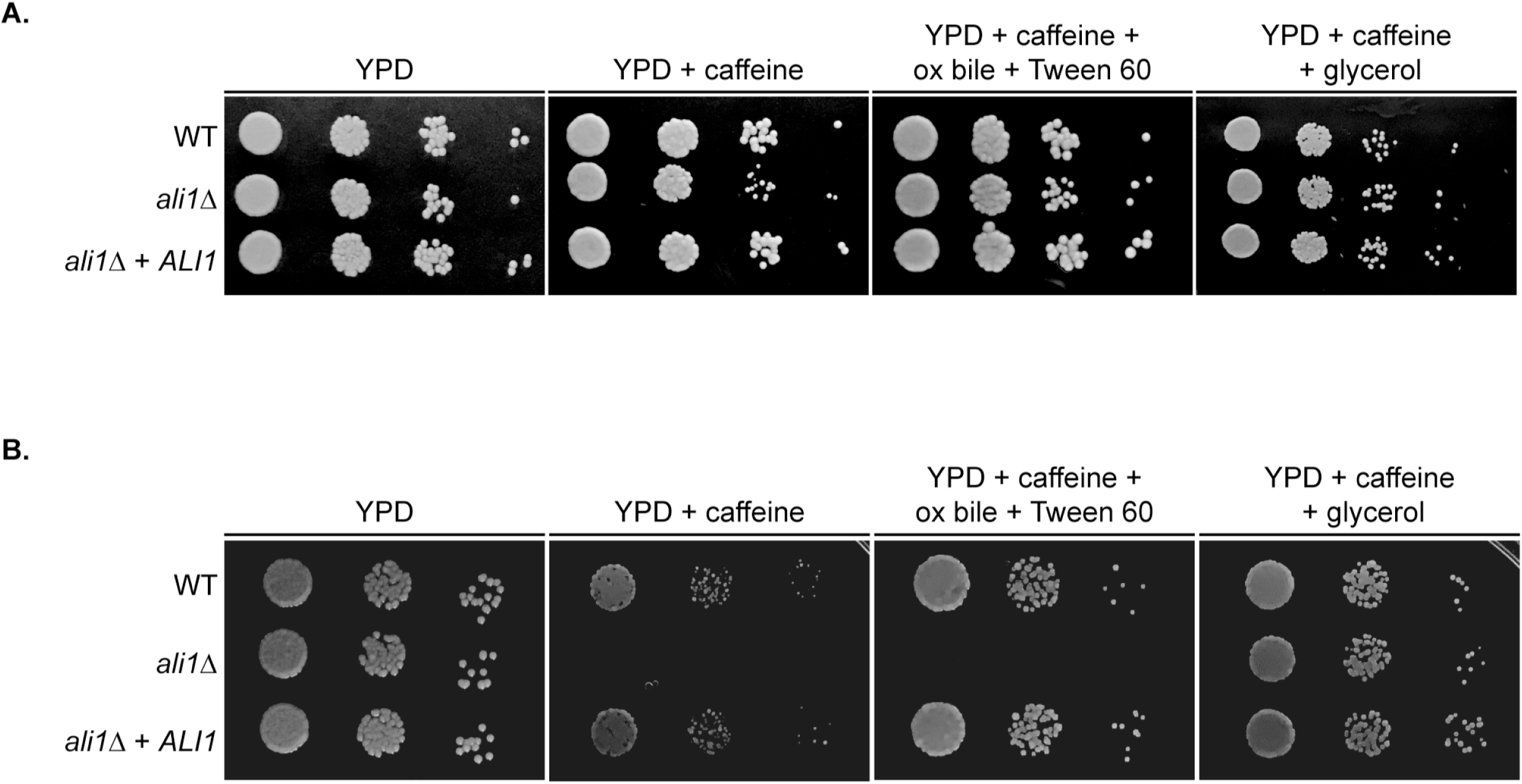
The effects of lipid supplementation on the *ali1*Δ mutant. Serial dilutions of the WT, *ali1*Δ mutant, and *ali1*Δ + *ALI1* strains were incubated on YPD medium; YPD with caffeine (1 mg/mL); YPD with caffeine, ox bile (10 mg/mL), and Tween 60 (1%); and YPD with caffeine and glycerol (0.4%). These strains were incubated at 30°C (A) and 37°C (B) and monitored visually for growth.

As well as its role as a cell surface stressor, caffeine is an inhibitor of the target of rapamycin complex 1 (TORC1) (36, 40). To determine if Ali1 functions in a pathway related to TORC1 function, we assessed the sensitivity of the *ali1*Δ mutant strain to rapamycin. We observed a two-fold decrease in the rapamycin MIC for the *ali1*Δ mutant compared to the WT strain at both 30°C (*ali1*Δ MIC_50_ = 1.56 ng/mL; WT MIC_50_ = 3.12 ng/mL) and 37°C (*ali1*Δ MIC_50_ = 0.78 ng/mL; WT MIC_50_ = 1.56 ng/mL). Inhibition of TORC1 induces autophagy in yeast (42). If caffeine-mediated inhibition of TORC1 is more effective in the *ali1*Δ mutant than in the WT strain, causing the *ali1*Δ mutant to display caffeine sensitivity, the *ali1*Δ mutant should be more susceptible to inducers of autophagy than the WT strain. Given the minimal change in rapamycin sensitivity of the *ali1*Δ mutant, we assessed the ability of the *ali1*Δ mutant strain to survive in nitrogen deprivation, a known inducer of autophagy (43). We incubated the WT strain, the *ali1*Δ mutant strain, and the *ali1*Δ + *ALI1* strain on synthetic low-ammonium dextrose (SLAD) medium at 30°C and 37°C. We observed that all strains displayed similar growth kinetics (data not shown). These data suggest that the caffeine sensitivity of the *ali1*Δ mutant is not due to dysregulation of autophagy, and that Ali1 likely does not directly function in a TORC1-related pathway.

In addition to the observation that the two strongest potential interactors of Ali1-GFP were Fas1 and Fas2, we also found that the fatty acid synthase inhibitor, cerulenin, is slightly more active against the *ali1*Δ mutant (MIC_50_ = 0.15 μg/mL) than the WT strain (MIC_50_ = 0.3 μg/mL). From these data, we hypothesized that the caffeine susceptibility of the *ali1*Δ mutant may be caused by impaired lipid synthesis. We supplemented the caffeine medium with various compounds involved in lipid synthesis and utilization, media additions that are frequently used to support the *in vitro* growth of lipid auxotrophic fungi such as *Malassezia* species (44). The addition of ox bile (10 mg/mL), which aids in the degradation and absorption of lipids, and Tween 60 (1%), which serves as an emulsifier, rescued the caffeine sensitivity of the *ali1*Δ mutant at 30°C, but not at 37°C (Fig. 4). The addition of glycerol (0.4%), a precursor for phospholipids and triglycerides, completely rescued the caffeine sensitivity of the *ali1*Δ mutant at both 30°C and 37°C (Fig. 4). In order to eliminate the possibility that glycerol was solely providing osmotic support that allowed for the *ali1*Δ mutant to overcome its caffeine sensitivity, we also supplemented the caffeine medium with sorbitol (1 M) and observed that it did not rescue the caffeine sensitivity of the *ali1*Δ mutant at either temperature (data not shown) (45, 46). Collectively, these observations indicate that lipid precursor supplementation is sufficient to suppress the caffeine sensitivity of the *ali1*Δ mutant, suggesting that the loss of cell surface integrity of the *ali1*Δ mutant is caused in part by impaired localized lipid synthesis and/or deposition, potentially at the site of cell separation.

### The *C. neoformans* arrestin family supports virulence *in vitro* and *in vivo*

Because we observed that the *ali1*Δ mutant exhibits phenotypes that are relevant to pathogenesis, specifically cytokinesis defects at elevated temperature and sensitivity to the cell surface stressor caffeine, we hypothesized that Ali1 may support fungal virulence. As a preliminary assessment, we evaluated the ability of the *ali1*Δ mutant to survive and proliferate in an *in vitro* macrophage co-culture system (26, 47, 48). We co-cultured the WT strain, the *ali1*Δ mutant, and the *ali1*Δ + *ALI1* strain for 24 hours with J774A.1 murine macrophages. We observed that the *ali1*Δ mutant displays a moderate, reproducible reduction in its ability to survive in the presence of macrophages compared to the WT strain, a phenotype that is rescued by complementation with the WT *ALI1* allele (Fig. 5A). We then performed *in vivo* studies in a murine inhalation model of cryptococcal infection (38, 48, 49). Following intranasal inoculation of C57BL/6 mice (n = 10) with 10^4^ colony forming units (CFU) of each strain, we observed no differences between the WT strain and the *ali1*Δ mutant in their abilities to cause lethal infection (Fig. 5B). From these results, we concluded that Ali1 has modest contributions to *in vitro* survival in the presence of macrophages, but does not promote *in vivo* virulence in a murine inhalation infection model.

**FIGURE 5.**
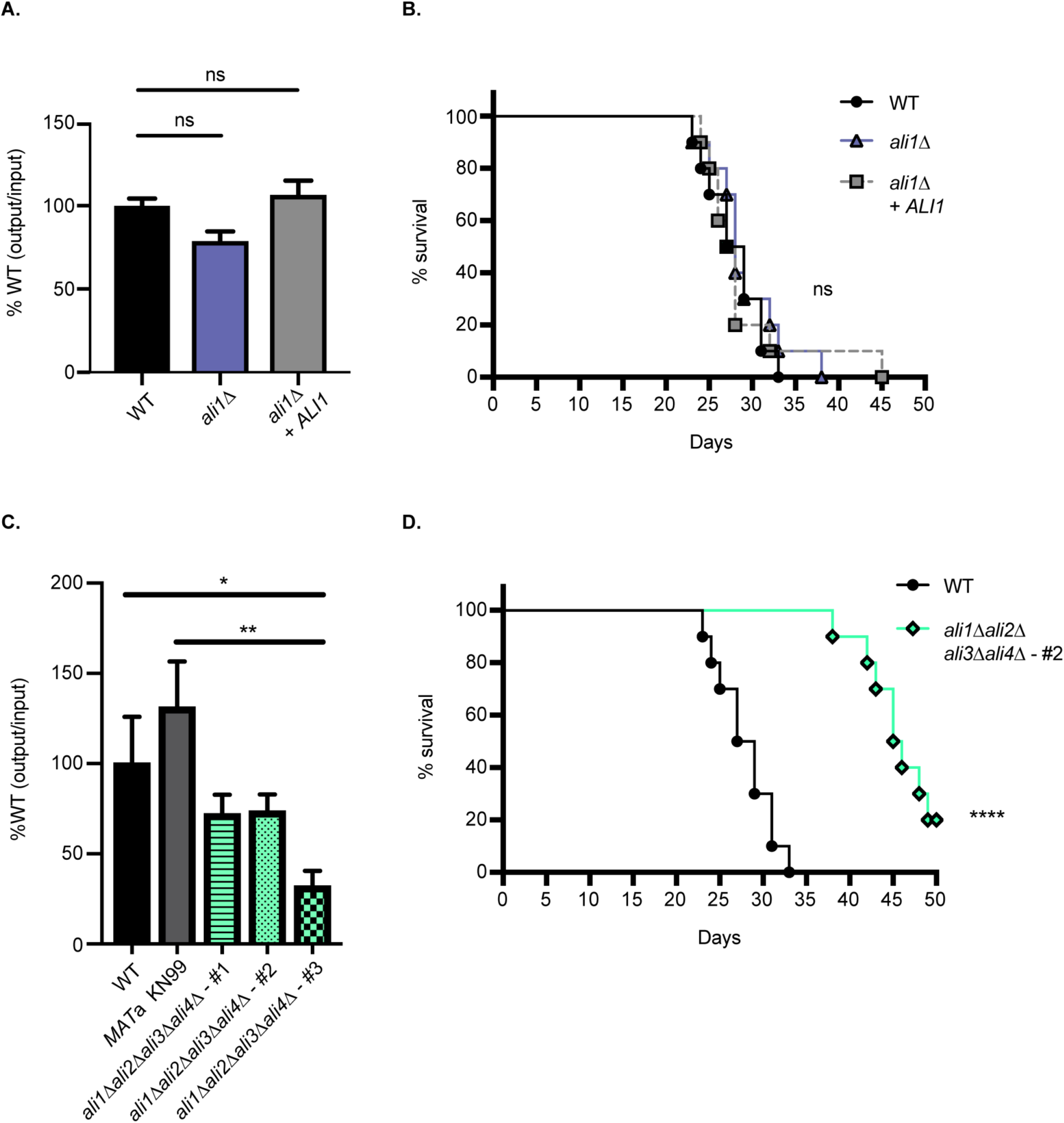
Virulence contributions of the *C. neoformans* arrestin family. A. The WT, *ali1*Δ mutant, and *ali1*Δ + *ALI1* strains were co-incubated with J774A.1 murine macrophages at a multiplicity of infection (MOI) = 1 for 24 hours. Survival of the strains was assessed by quantitative culture, and the percentage of recovered colony-forming units (CFU) was normalized to the WT strain. This experiment was performed with six biological replicates (n = 6). Error bars represent the SEM. Log transformation was used to normally distribute the data for statistical analysis (One-way ANOVA; ns = not significant). B. Female C57BL/6 mice (n = 10) were intranasally inoculated with 10^4^ CFU of the WT, *ali1*Δ mutant, or *ali1*Δ + *ALI1* strains. Mouse survival was tracked for 50 days post-infection (Log-rank test; ns = not significant). C. The WT strain, the *MAT*a KN99 strain, and three isogenic but independent arrestin null mutants (all also *MAT*a) were co-incubated with J774A.1 murine macrophages at a MOI = 1 for 24 hours. Survival of the strains was assessed by quantitative culture, and the percentage of recovered CFU was normalized to the WT strain. This experiment was performed with four biological replicates (n = 4). Error bars represent the SEM. Log transformation was used to normally distribute the data for statistical analysis (*One-way ANOVA p < 0.05; **One-way ANOVA p < 0.01). D. Female C57BL/6 mice (n = 10) were intranasally inoculated with 10^4^ CFU of the WT strain and a representative arrestin null mutant, *ali1*Δ*ali2*Δ*ali3*Δ*ali4*Δ - #2 (CLT57). Mouse survival was tracked for 50 days post-infection (****Log-rank test p < 0.0001).

Because the *ali1*Δ mutant individually does not exhibit significant virulence defects, we next determined whether the *C. neoformans* arrestin family, collectively, contributes to virulence. To do so, we utilized the *ali1*Δ*ali2*Δ*ali3*Δ*ali4*Δ mutants, referred to as the “arrestin null” mutants, in which all four known *C. neoformans* arrestins are ablated. Similar to our studies with the *ali1*Δ mutant, we evaluated the ability of three independent arrestin null mutants to survive and proliferate in an *in vitro* macrophage co-culture system (26, 47, 48). To do so, we co-cultured the WT strain, the *MAT*a KN99 strain (which was used in genetic crosses to generate the arrestin null mutants), and three arrestin null mutants for 24 hours with J774A.1 murine macrophages. We observed that all three arrestin null mutants exhibit a marked reduction in their abilities to survive in the presence of macrophages, compared to the WT strain and the *MAT*a KN99 strain (Fig. 5C). A representative arrestin null mutant, *ali1*Δ*ali2*Δ*ali3*Δ*ali4*Δ - #2 (CLT57), was then assessed for virulence in the murine inhalation model (38, 48, 49). Following intranasal inoculation of C57BL/6 mice (n = 10) with 10^4^ CFU of the WT strain or the arrestin null mutant, we observed that the arrestin null mutant displays a significant attenuation in virulence compared to the WT strain (Fig. 5D). Mice infected with the WT strain exhibited a median survival time of 28 days, while those infected with the arrestin null mutant exhibited a median survival time of 45.5 days (Fig. 5D). These data collectively indicate that the *C. neoformans* arrestin family contributes to both *in vitro* and *in vivo* virulence.

### The *C. neoformans* arrestins likely serve distinct cellular functions

In order to identify possible mechanisms by which the *C. neoformans* arrestin family contributes to virulence, we created individual *ali1*Δ, *ali2*Δ, *ali3*Δ, and *ali4*Δ loss-of-function mutants. Following strain confirmation, we assessed the growth kinetics of the arrestin mutants in the presence of various cellular stressors. Specifically, we incubated the WT strain, the individual arrestin mutants, and the arrestin null mutants in the presence of physiologically-relevant stressors, such as elevated temperature (39°C), high salt (1.5 M NaCl), and alkaline pH (pH 8), as well as cell surface stressors, such as caffeine (1 mg/mL) and SDS (0.03%) (25, 39, 40, 50). We observed that the individual arrestin mutants display distinct, but overlapping, phenotypes in the presence of these stressors (Fig. 6). All of these individual arrestin mutant phenotypes are rescued by complementation with the respective WT arrestin allele (Fig. S2).

**FIGURE 6.**
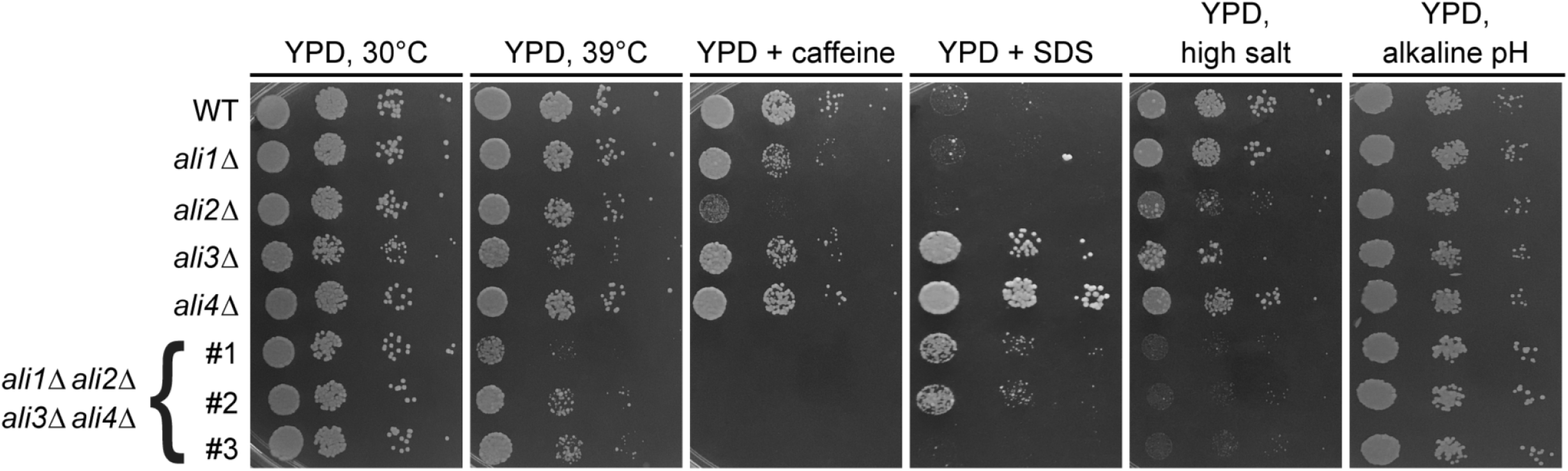
The *C. neoformans* arrestin mutant phenotypes. Serial dilutions of the WT strain, each individual arrestin mutant, and three independent arrestin null mutants were incubated on YPD medium with the following growth conditions/additives: 30°C, 39°C, caffeine (1 mg/mL), SDS (0.03%), high salt (1.5 M NaCl), and alkaline pH (pH 8). Cells were monitored visually for growth. The *ali1*Δ mutant exhibits modest susceptibility to caffeine. This phenotype is shared by, but markedly enhanced in, the *ali2*Δ mutant. The *ali2*Δ mutant also displays sensitivity to high salt. The *ali3*Δ mutant has modest growth defects at 39°C, as well as resistance to SDS. The *ali4*Δ mutant shares this SDS resistance phenotype, although it is enhanced compared to the *ali3*Δ mutant. The arrestin null mutants display reduced growth in the presence of 39°C, caffeine, and salt, but enhanced growth in the presence of SDS and alkaline pH.

Because we observed that the *ali2*Δ mutant has an enhanced caffeine sensitivity phenotype compared to the *ali1*Δ mutant, we hypothesized that the *ali2*Δ mutant would display more severe virulence defects than the *ali1*Δ mutant. To test this hypothesis, we co-cultured the WT strain, the *ali2*Δ mutant, the *ali2*Δ + *ALI2-GFP* strain, and an *ali1*Δ*ali2*Δ mutant for 24 hours with J774A.1 murine macrophages. The *ali2*Δ mutant had a significant reduction in its ability to survive in the presence of macrophages compared to the WT strain, a phenotype that is rescued by complementation with the WT *ALI2* allele (Fig. S3). The *ali2*Δ mutant survival rate (67%) is lower than what was observed for the *ali1*Δ mutant (79%) (Fig. 5A). Additionally, the *ali1*Δ*ali2*Δ mutant exhibits a more severe survival impairment (43%) than either the *ali1*Δ mutant or *ali2*Δ mutant alone, indicating that Ali1 and Ali2 have additive effects that contribute to survival in the presence of macrophages.

The arrestin null mutants share many phenotypes with the individual arrestin mutants, such as sensitivity to high temperature, caffeine, and high salt, as well as resistance to SDS (Fig. 6). Uniquely, the arrestin null mutants display a slight increase in growth rate in the presence of alkaline pH (Fig. 6). The most pronounced phenotypes of the arrestin null mutants, growth defects in the presence of high temperature and caffeine, were not rescued by glycerol (0.4%) supplementation but were partially rescued by osmotic support with sorbitol (1M) supplementation (Fig. S4). These data suggest that the *C. neoformans* arrestin proteins likely perform distinct, nonredundant cellular functions that contribute to survival in physiologically-relevant conditions and cell surface stability.

## DISCUSSION

### Arrestins have been well-characterized in model fungi systems

The model ascomycete fungi, such as *S. cerevisiae*, *Aspergillus nidulans*, and *Schizosaccharomyces pombe*, all contain relatively large α-arrestin families of nine to eleven members (29). Based on the presence of the conserved arrestin domains, α-arrestins are predicted to exist in the other three major fungal groups: the basidiomycetes, the zygomycetes, and the chytrids (29). We used *C. neoformans* as a genetically-tractable basidiomycete, with a relatively small arrestin family of four members, to more broadly characterize fungal α-arrestin functions, both individually and collectively. Additionally, because *C. neoformans* is a major human pathogen, we interrogated the functional contributions of fungal α-arrestins to virulence. The fact that the α-arrestins, despite lacking catalytic activity themselves, have remained present within all major fungal groups indicates that they are likely functionally important proteins within the fungal kingdom.

### Ali1 is important for cytokinesis in the presence of cellular stress

Septins are conserved GTP-binding proteins that create the septum in eukaryotes, often serving as scaffolds for other proteins that direct cell cycle progression (51–53). In *S. cerevisiae*, the septin proteins assemble into filaments at the mother bud neck, creating the hourglass-shaped septum, and are required for normal cytokinesis (53). The *C. neoformans* septins have been shown to function similarly. *C. neoformans* septin mutants display cytokinesis defects when incubated at elevated temperatures and also display modest sensitivity to cell surface stressors, such as caffeine and SDS (34).

We observed that Ali1 has cell cycle-associated localization, with enrichment at the septum and poles of budding cells. Our protein interactome analysis supported this observation, with multiple septin proteins, Cdc10, Cdc11, and Cdc12, identified at low levels in the Ali1-GFP immunoprecipitations. Protein-protein interactions with septin proteins are typically transient, potentially explaining the low APC for the septin proteins using this experimental approach (54). Additionally, we found that the *ali1*Δ mutant displays an increased incidence of cytokinesis defects at elevated temperature and sensitivity to the cell surface stressor caffeine, thus phenocopying the *C. neoformans* septin mutants (34). These data suggest that Ali1 is a regulator of cytokinesis that is particularly important in the presence of stress. Whole transcriptome analyses of synchronized *C. neoformans* cells have shown that Ali1 expression is cyclic, or regulated with the cell cycle, with its peak expression occuring about 15 minutes prior to bud emergence (55). As a potential regulator of cytokinesis, this expression pattern would enable the *ALI1* transcript to be transcribed, and the Ali1 protein to be translated and localize to the septum and poles as cell division is occuring.

In addition to and in collaboration with septins, Ras GTPases are conserved regulators of cell division in eukaryotes. Our laboratory has shown that the *C. neoformans* Ras1 protein directs polarized growth and actin polarization, particular in the presence of stress (22, 23, 33). When Ras1 is inhibited, septins are unable to organize at the septum to perform their scaffolding functions and cells display morphological and cytokinesis defects (23). We demonstrated that in the absence of Ras1, Ali1 localization to the septum and poles is impaired. This observation indicates that the cell cycle-regulated localization of Ali1 is dependent on Ras1.

### Ali1 likely fulfills an adaptor role aiding cytokinesis

Cytokinesis is a highly organized and regulated process in fungi. In *S. cerevisiae*, cell wall enzymes, such as the β(1–3)-glucan synthases and the chitin synthases, localize to the septum and poles to help build the septum and cell wall during cell division. (56–58). It is believed that *C. neoformans* also directs cytokinesis similarly. For example, *C. neoformans* cells lacking Chs3, a chitin synthase, or Ags1, the α(1–3)-glucan synthase, display cytokinesis defects during budding (59, 60). Similar to the cell wall, the cell membrane must be remodeled to aid in bud growth and cytokinesis in fungi. To our knowledge, little work has focused on the degradation and rebuilding of the fungal cell membrane during cytokinesis. However, in the bacterium *Mycobacterium tuberculosis*, fatty acid synthase proteins localize to the poles and septum to synthesize the mycomembrane during cell division (61).

Fungal fatty acid synthases, which belong to the microbial type I fatty acid synthase family, are cytosolic multi-enzymes that heterodimerize to form hexamers (α6β6) (62–64). Once in this complex, they employ their individual component enzymes to synthesize de novo a diversity of lipid products that are used for cellular metabolism, signaling, and biological membranes. In *C. neoformans*, Fas1 and Fas2 are required for viability in standard laboratory conditions and are targets of the fatty acid synthase inhibitor cerulenin (65). Through our protein interactome analysis, we found that the two strongest potential interactors of Ali1 are Fas1 and Fas2. We tested the sensitivity of the *ali1*Δ mutant to cerulenin and observed that the *ali1*Δ mutant strain is slightly more sensitive to cerulenin than the WT strain. In conjunction with these data, we observed that the *ali1*Δ mutant displays sensitivity to the cell surface stressor caffeine, which is enhanced in the presence of temperature stress. In addition to its roles as a cell surface stressor, caffeine is believed to inhibit TORC1 (36, 40). The caffeine sensitivity of the *ali1*Δ mutant may be explained by the fact that TORC1 is an upstream activator of lipid synthesis genes in eukaryotes, including fatty acid synthases (66–68). Supplementation with exogenous lipid precursors, but not the osmotic stabilizer sorbitol, may reverse the caffeine sensitivity of the *ali1*Δ mutant by compensating for an insufficiency in substrates used to synthesize cellular membranes. These data collectively suggest that Ali1 is required for complete Fas1 and Fas2 function.

The *S. pombe* α-arrestin, Art1, regulates cytokinesis through its adaptor function (69). Art1 is required for the localization of Rgf3, the guanine nucleotide exchange factor for the regulatory subunit of the β-glucan synthase, Rho1, to the septum, likely so that it can help build the septum. Our data suggest that Ali1 functions similarly to Art1. We hypothesize that Ali1 acts as an adaptor for Fas1 and Fas2, aiding in their localization to the septum and poles, so that they can rebuild the cell membrane during cytokinesis (Fig. 7). In the absence of Ali1, cells are left with small, localized defects in the cell surface because they are unable to repair the membrane, or are delayed in membrane repair, compared to WT cells, particularly in the presence of stress. This results in the cytokinesis and cell surface defects observed in the *ali1*Δ mutant. Previous work in both mammals and fungi have demonstrated the importance of fatty acid synthesis for progression through the cell cycle (70–72). Additionally, the mechanism by which Ali1 is able to perform its adaptor function for Fas1 and Fas2 may be ubiquitin-mediated, through interactions with the E3 ubiquitin ligase Rsp5 (Fig. 7). Ali1 contains four potential ubiquitin ligase binding sites, or PxY sites. We also observed that Ali1 interacts with multiple ubiquitination proteins, including E1, E2, and E3 proteins, particularly in TC conditions (73). Ubiquitination is most often considered in the context of proteasomal degradation, but it can also direct diverse subcellular localizations (74–76).

**FIGURE 7.**
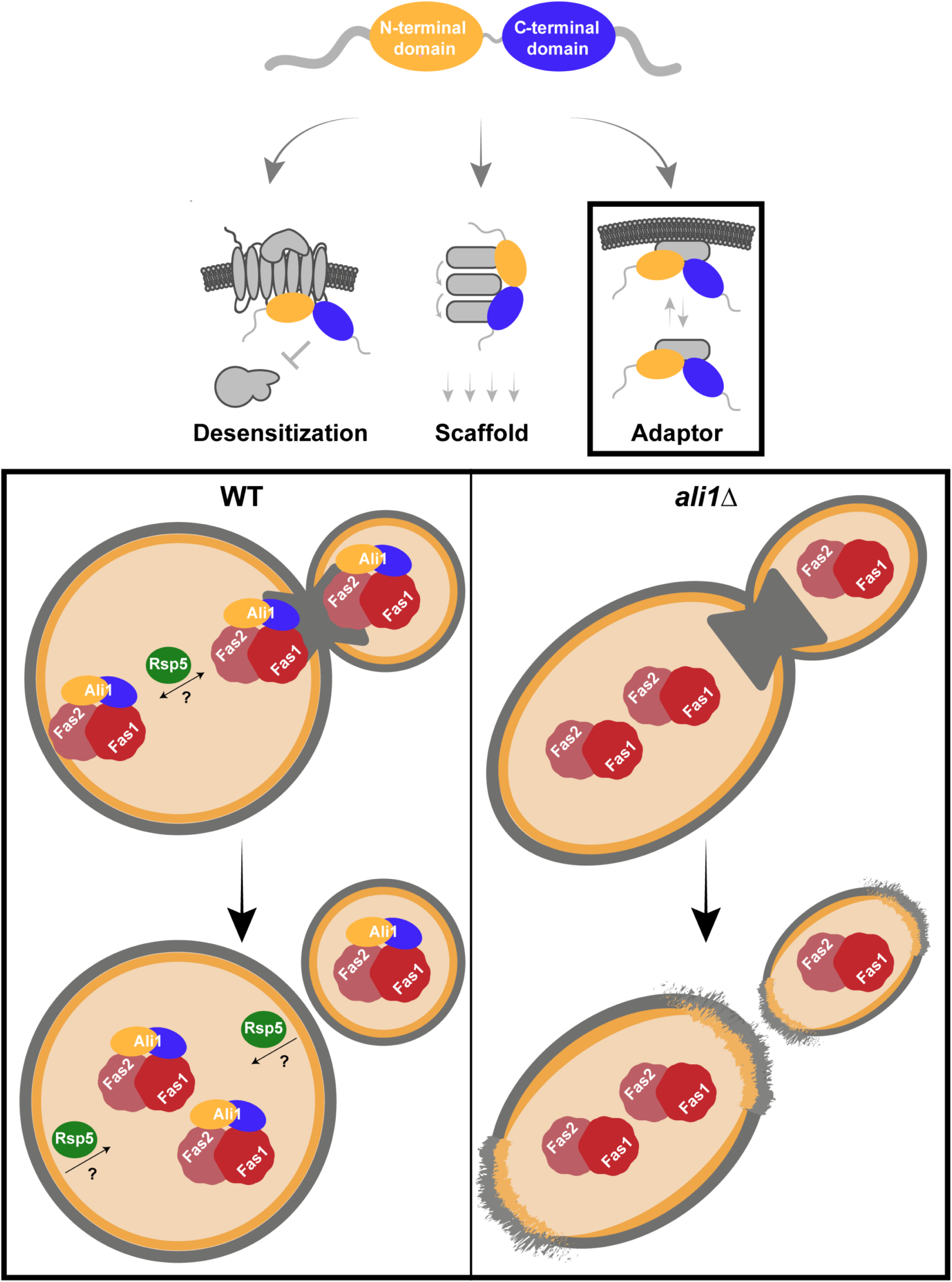
Working model of Ali1 adaptor function. We propose that Ali1 (yellow/blue) acts as an adaptor protein to aid in the localization of the Fas1 and Fas2 fatty acid synthase complex (red) to the septum and poles of actively dividing cells, possibly in a ubiquitin-mediated manner through interactions with the E3 ubiquitin ligase Rsp5 (green). This process occurs to help meet the increased, stress-induced need for lipid synthesis and deposition at these sites. In the *ali1*Δ mutant strain, Fas1 and Fas2 are unable to localize, or are delayed in their localization, to the septum and poles during cell division. As a result, lipid synthesis and deposition at these sites is impaired. This causes localized cell surface defects at the poles in the resulting cells, likely explaining the cytokinesis defects and caffeine sensitivity phenotypes of the *ali1*Δ mutant.

Future investigations can explore the interactions between Ali1 and the fatty acid synthases, Fas1 and Fas2. Additionally, the localizations of Fas1 and Fas2 in the WT and the *ali1*Δ mutant backgrounds can also be assessed. However, it is possible that it may be difficult to draw conclusions from these experiments. Fas1 and Fas2 are abundant, diffusely cytosolic proteins in *S. cerevisiae* (77). If this is also the case for *C. neoformans*, it may be challenging to observe any transient interactions or enrichments of these proteins at the septum and poles.

### The *C. neoformans* arrestin family contributes to virulence

Upon infection, pathogens must regulate their adaptive cellular responses to acclimate to the stressors of the host environment. Work largely conducted in ascomycete fungi has demonstrated that disruption of α-arrestin-regulated adaptive cellular responses is detrimental for fungal survival and pathogenesis. For example, the α-arrestin Rim8 scaffolds the Rim alkaline pH response pathway in *Candida albicans*; the *rim8*Δ mutant displays attenuation in a murine model of systemic candidiasis, indicating that Rim8 is required for adaptation to the host environment (78). Given many investigations demonstrating that human arrestin proteins regulate cellular processes that are involved in human disease, we propose that fungal arrestins similarly regulate fungal adaptive cellular responses important for disease establishment and progression (79–82).

This study directly investigates the virulence contributions of fungal α-arrestins. Implementing a murine inhalation model of cryptococcal infection, we observed that the individual *ali1*Δ mutant does not display virulence attenuation, but that the arrestin null mutant exhibits a significant delay in its ability to cause fatal disease. These data suggest that the arrestins, collectively, are involved in adaptation to the host environment in *C. neoformans*. Since we observed that the *ali2*Δ mutant displays more severe attenuation in its ability to survive in the presence of macrophages than the *ali1*Δ mutant, we propose that Ali2 is a compelling subject for future investigations. Additionally, because the arrestin mutants have distinct phenotypes in the presence of different cellular stressors, as well as because the *C. neoformans* arrestin family is very small, we hypothesize that the *C. neoformans* arrestins have distinct cellular functions that contribute to adaptation to the host. Functional redundancy has been observed for mammalian and fungal arrestins, therefore it is possible that the *C. neoformans* arrestins could have some degree of overlapping functions while maintaining protein-specific activities as well (19, 83).

We have demonstrated that the *C. neoformans* arrestin family contains four members that share little primary amino acid sequence conservation with human arrestins. These fungal-specific proteins likely mediate various cellular functions including efficient progression through the cell cycle, especially under stressful growth conditions. Fungal arrestins therefore offer unique insight into mechanisms of stress response and cellular adaptation in this diverse group of eukaryotes.

## MATERIALS AND METHODS

### Strains, media, and growth conditions

All strains used in this study were generated in the *C. neoformans* var. *grubii* H99 (*MAT*α) or KN99 (*MAT*a) backgrounds and are included in Table 3. Strains were maintained on yeast extract-peptone-dextrose (YPD) medium (1% yeast extract, 2% peptone, 2% dextrose, and 2% agar for solid medium). To regulate *RAS1* expression, yeast extract-peptone-galactose (YPGal) medium (1% yeast extract, 2% peptone, and 3% galactose) was utilized (23). CO_2_-independent tissue culture (TC, Gibco) medium was used to mimic an *in vivo* environment, as described previously (84). To assess mutant strain cell surface phenotypes, NaCl (1.5 M) and Congo red (0.5%) were added to YPD medium before autoclaving, while caffeine (1 mg/mL), calcofluor white (1 mg/mL), and SDS (0.03%) were filter sterilized and added to YPD medium after autoclaving (38). Synthetic low-ammonium dextrose (SLAD) medium (0.17% yeast nitrogen base without amino acids and without ammonium sulfate, 50 μM ammonium sulfate, 2% dextrose, and 2% agar) was used as a nitrogen deprivation medium to induce autophagy. Lipid precursor supplementation was achieved by adding ox bile (HiMedia Labs) (10 mg/mL) and Tween 60 (1%) to medium before autoclaving, or by adding sterile glycerol (0.4%) to medium after autoclaving. Sorbitol supplementation was achieved by adding sorbitol (1M) to medium before autoclaving. Alkaline pH plates were made by adding 150 mM HEPES buffer to YPD medium and adjusting the pH to 8.15 with NaOH prior to autoclaving (25). Unless otherwise indicated, strains were incubated at 30°C.

**TABLE 3.**
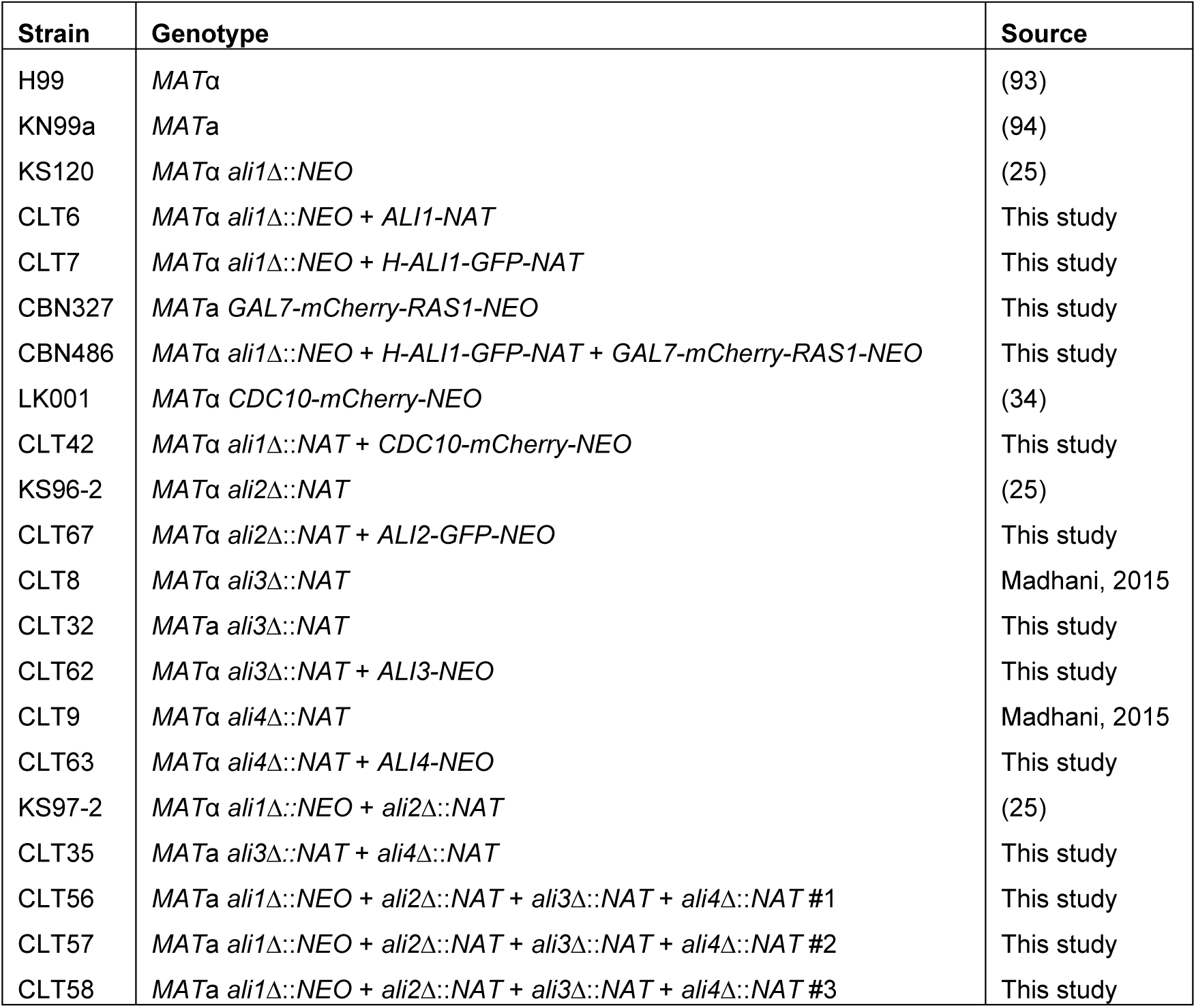
Strains used in this study

### Molecular biology and strain construction

All plasmids used in this study are listed in Table 4. All primers utilized in this study are listed in Table 5. All strains were generated by biolistic transformation, unless otherwise described (85). Detailed methods for the construction of all strains used in this study are included in File S1 (86–88).

**TABLE 4.**
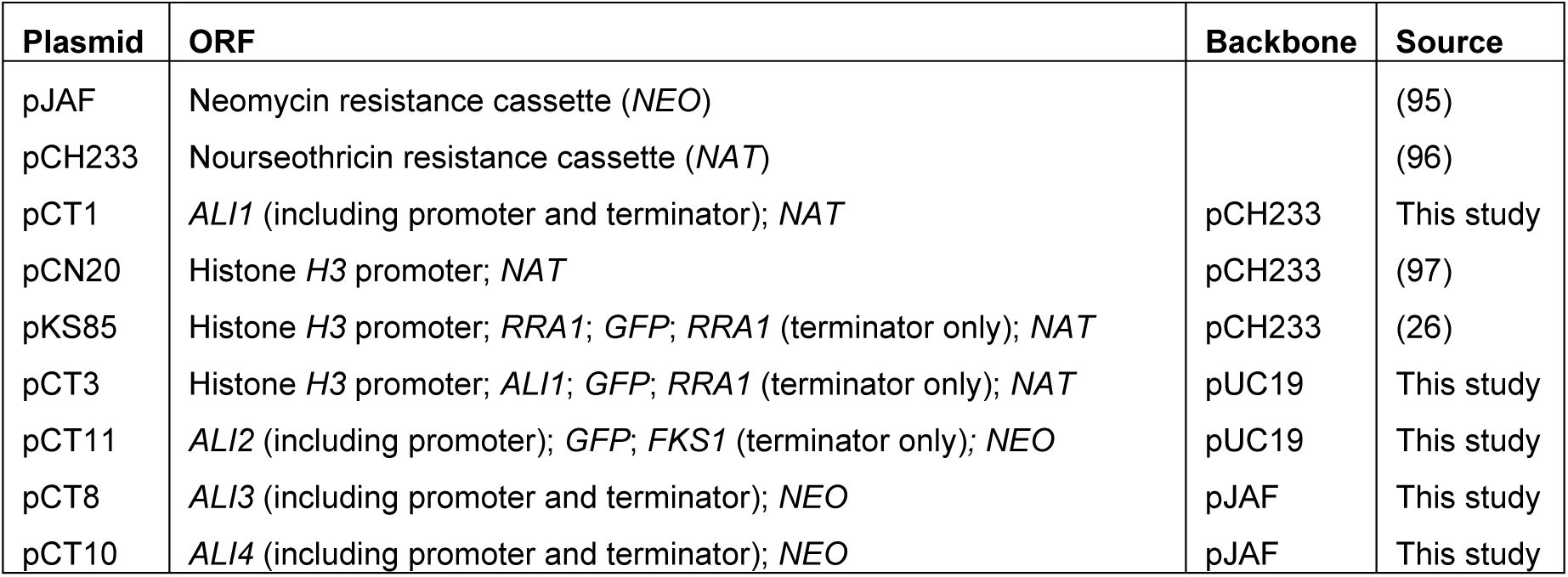
Plasmids used in this study

| Plasmid | ORF | Backbone | Source |
| --- | --- | --- | --- |
| pJAF | Neomycin resistance cassette (*NEO*) |  | (95) |
| pCH233 | Nourseothricin resistance cassette (*NAT*) |  | (96) |
| pCT1 | *ALI1* (including promoter and terminator); *NAT* | pCH233 | This study |
| pCN20 | Histone *H3* promoter; *NAT* | pCH233 | (97) |
| pKS85 | Histone *H3* promoter; *RRA1*; *GFP*; *RRA1* (terminator only); *NAT* | pCH233 | (26) |
| pCT3 | Histone *H3* promoter; *ALI1*; *GFP*; *RRA1* (terminator only); *NAT* | pUC19 | This study |
| pCT11 | *ALI2* (including promoter); *GFP*; *FKS1* (terminator only); *NEO* | pUC19 | This study |
| pCT8 | *ALI3* (including promoter and terminator); *NEO* | pJAF | This study |
| pCT10 | *ALI4* (including promoter and terminator); *NEO* | pJAF | This study |

**TABLE 5.**
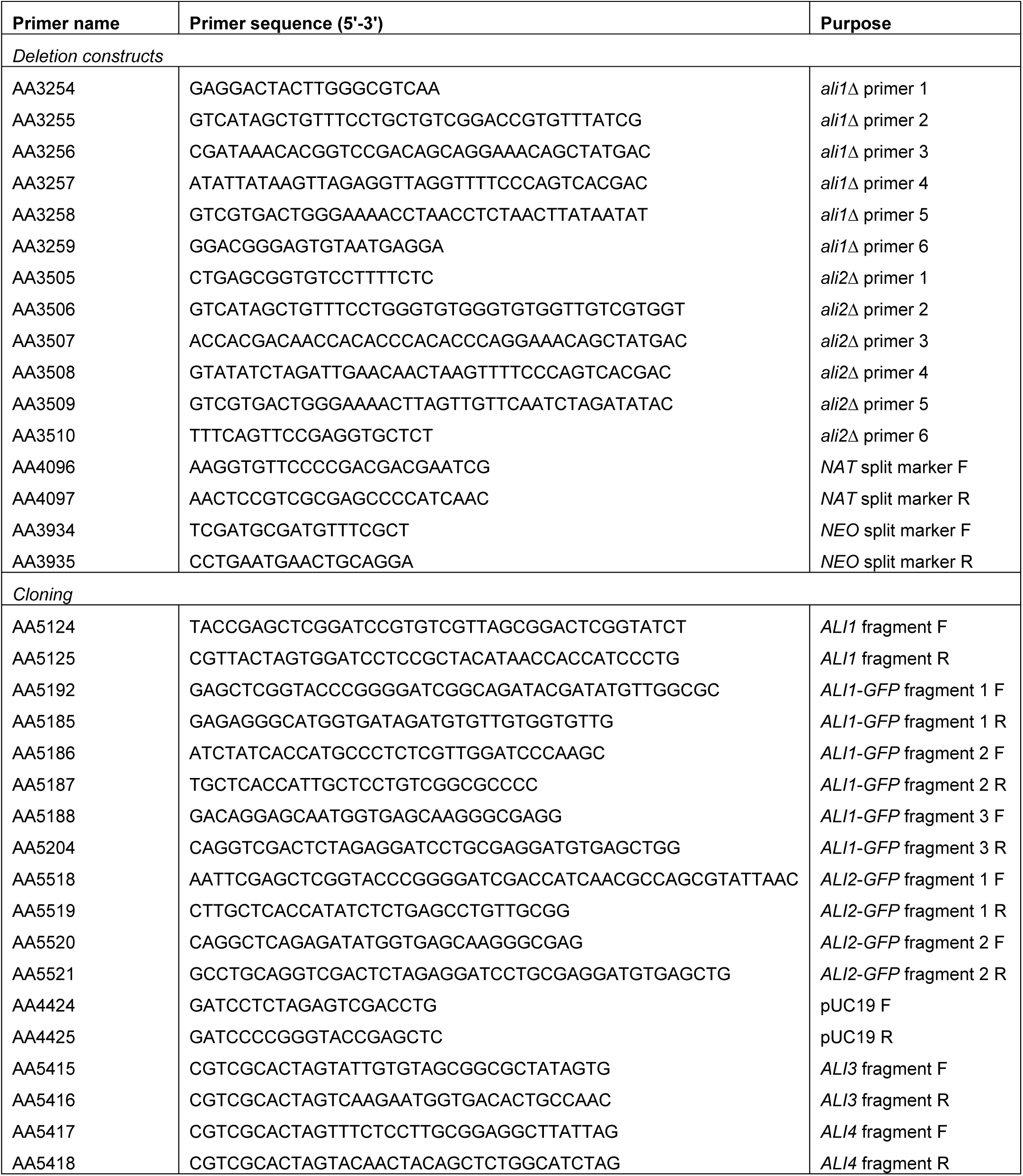
Primers used in this study

### BLAST analyses

To identify homology between the *C. neoformans* arrestins and those in *S. cerevisiae* and humans, Basic Local Alignment Search Tool (BLAST, NCBI) was used. The protein sequences of each of the *C. neoformans* arrestins was searched against the *S. cerevisiae* S288C (taxid:559292) and human (taxid:9606) proteomes using the default parameters for protein-protein BLAST (blastp) and Position-Specific Iterated BLAST (PSI-BLAST) (30–32). Alignments considered significant, those with E values less than 1, are included in Tables S1 (*S. cerevisiae*) and S2 (human).

### Fluorescent and light microscopy

All images in this study (differential interference contrast [DIC] and fluorescent) were captured using a Zeiss Axio Imager A1 microscope equipped with an Axio-Cam MRM digital camera. To assess subcellular localization of Ali1-GFP, the WT (H99) strain and the Ali1-GFP (CLT7) strain were incubated for 18 hours with 150 rpm shaking in YPD medium at 30°C or TC medium at 37°C. Cells were then pelleted, washed with phosphate-buffered saline (PBS), and imaged.

To measure the frequency of cell cycle-associated localization of Ali1-GFP in the presence and absence of Ras1, the Ali1-GFP + mCherry-Ras1 (CBN486) strain was incubated for 18 hours at 30°C with 150 rpm shaking in YPGal medium. Cells were pelleted, washed three times with PBS, normalized by spectrophotometry, and then resuspended to an OD_600_ of 0.2 in either YPGal medium (to induce *RAS1* expression) or YPD medium (to repress *RAS1* expression) for 18 hours at 30°C with 150 rpm shaking (23). Cells were then pelleted, washed with PBS, and imaged. Results are reported as the average percentage (+/-standard error of the mean [SEM]) of actively budding cells that displayed Ali1-GFP localization to the septum and/or poles. Statistical significance was determined using Student’s *t*-test (GraphPad Software, San Diego, CA). A minimum of 600 cells were analyzed in both YPGal and YPD conditions across three biological replicates using ImageJ Software (Fiji) (89, 90).

To analyze the morphology of the *ali1*Δ mutant cells, the WT (H99), *ali1*Δ (KS120), and *ali1*Δ + *ALI1* (CLT6) strains were incubated for 18 hours at 30°C with 150 rpm shaking in YPD medium. An OD of approximately 0.2 for each strain was transferred to fresh YPD medium and subsequently incubated at either 30°C or 39°C for 18 hours with 150 rpm shaking. Cells were then pelleted, washed with PBS, and imaged. Results are reported as the average percentage (+/-SEM) of total cells displaying morphological defects. Statistical significance was determined using one-way analysis of variance (ANOVA) and the Tukey-Kramer test (GraphPad Software, San Diego, CA). A minimum of 600 cells were analyzed across three biological replicates using ImageJ Software (Fiji) (89, 90).

To assess whether Ali1 is required for septin protein localization, the Cdc10-mCherry (LK001) strain and the Cdc10-mCherry + *ali1*Δ (CLT42) strain were incubated for 18 hours at either 30°C or 37°C with 150 rpm shaking in YPD medium. Cells were then pelleted, washed with PBS, and imaged.

### Protein isolation, membrane fractionation, and western blotting

For all protein experiments, protein extracts were prepared as previously described (25, 26). Briefly, the WT (H99) and the Ali1-GFP (CLT7) strains were incubated for 18 hours at 30°C with 150 rpm shaking in YPD medium. Cells were pelleted, flash frozen on dry ice, and lysed by bead beating. The crude lysate was cleared by centrifugation at 2,500 x g at 4°C for 5 minutes and the supernatant (total cell lysate) was transferred to a new tube. Total cell lysate protein concentrations were measured using bicinchoninic acid assay (BCA).

To determine the relative abundance of Ali1 in different cellular fractions, WT (H99) and the Ali1-GFP (CLT7) strains were incubated and lysed as above. Total cell lysates (T) were separated by ultracentrifugation at 30,000 x g for 1 hour at 4°C (27). The soluble fraction (S) was transferred to a new tube and the insoluble pellet (I) was resuspended in an equivalent volume of lysis buffer containing 1% Triton X-100. All samples were normalized by total protein concentration. Western blots were performed as described previously using an anti-GFP primary antibody (1/5,000 dilution, Roche) followed by an anti-mouse peroxidase-conjugated secondary antibody (1/25,000 dilution, Jackson Labs). Proteins were detected by enhanced chemiluminescence (ECL Prime Western blotting detection reagent, GE Healthcare).

### Proteomic experiment preparation and analysis

Proteomic analysis was performed with a single replicate for the WT (H99) strain in both YPD and TC conditions, and in triplicate for the Ali1-GFP (CLT7) strain in both YPD and TC conditions. To prepare total cell lysates for this experiment, the WT (H99) strain and the Ali1-GFP (CLT7) strain were incubated for 18 hours at 30°C with 150 rpm shaking in YPD medium. Both strains were normalized to an OD_600_ of 1, resuspended in either YPD or TC media, and then incubated for 3 hours at 30°C with 150 rpm. Cells were pelleted and lysed as described above to extract total cell lysates. Immunoprecipitations from total cell lysates were performed by addition of 25 µL GFP-Trap resin (Chromotek) and inversion at 4°C for 2 hrs. Mass spectrometry analysis was performed on immunoprecipitations by the Duke Proteomics Core Facility, as described previously (26). A description of this analysis is included in File S1.

We prioritized hits from this proteomic analysis to enrich for proteins with stronger potential interactions with Ali1-GFP. First, we averaged the exclusive unique peptide counts (APC) for each potential interactor identified in YPD or TC conditions, and subsequently selected those that had an APC of 2 or more for further analysis. We then calculated the percentage of the APC that was identified in the respective WT immunoprecipitation for each potential protein interactor. Those proteins that had less than 20% of the APC identified in the respective WT immunoprecipitation were determined to be unique interactors of Ali1-GFP. All proteins identified using this prioritization scheme in YPD and TC conditions can be found in Tables S4 and S5, respectively. All 1,122 identified proteins, except for those not belonging to *C. neoformans*, are included in Table S3.

### Macrophage co-culture experiments

The ability of strains to survive in the presence of macrophages was assessed as previously described (26). Briefly, 10^5^ J774.1 macrophages were incubated with 10^5^ opsonized fungal cells – WT (H99), KN99a, *ali1*Δ (KS120), *ali1*Δ + *ALI1* (CLT6), *ali2*Δ (KS96-2), *ali2*Δ + *ALI2-GFP* (CLT67), *ali1*Δ*ali2*Δ (KS97-2), and the arrestin null (CLT56, CLT57, and CLT58) mutants. Co-cultures of J774.1 macrophages and phagocytosed fungal cells were incubated for 24 hours at 37°C with 5% CO_2_. Phagocytosed fungal cells were collected, serially diluted, and plated onto YPD agar to assess the number of viable *C. neoformans* cells by quantitative culture. Results are reported as the average percentage (+/-SEM) of recovered colony-forming units (CFU), normalized to the WT (H99) strain, generated from at least 4 biological replicates. Statistical significance was determined using one-way analysis of variance (ANOVA) and the Tukey-Kramer test (GraphPad Software, San Diego, CA).

### Mouse survival experiments

The murine inhalation model of cryptococcosis was used to assess virulence of the stains in this study (49). C57BL/6 female mice were acquired from Charles Rivers Laboratories. Mice were anesthetized with 2% isoflurane utilizing a rodent anesthesia device (Eagle Eye Anesthesia, Jacksonville, FL) and were infected via the intranasal route with 10^4^ CFU of either the WT (H99) strain, the *ali1*Δ (KS120) mutant, the *ali1*Δ + *ALI1* (CLT6) strain, or an arrestin null (CLT57) mutant in 30 μl of sterile PBS. Mice (n = 10) were monitored twice daily and sacrificed if moribund. Survival data were statistically analyzed using log-rank test (GraphPad Software, San Diego, CA). Animal experiments were approved by The University of Texas at San Antonio Institutional Animal Care and Use Committee (IACUC) and mice were handled according to IACUC guidelines.

### Minimum inhibitory concentration (MIC) testing

To measure strain susceptibilities to rapamycin and cerulenin, MIC testing was performed using species-specific modifications to standard CLSI testing methods for broth microdilution testing of antifungal susceptibility (91, 92). A detailed description of this method is described in File S1.

## ACKNOWLEDGEMENTS

This work was supported by NIH R01 grant AI074677 (J.A.A.). We would like to thank the Duke Proteomics Core Facility for assistance with our proteomic experiments. We would also like to acknowledge the Madhani laboratory at UCSF and NIH funding R01AI100272 for the publicly available deletion mutant collections in *C. neoformans* (Fungal Genetics Stock Center, 2015/2016 Madhani plates).

---

**TABLE S1.** Primary amino acid sequence homology between the *C. neoformans* arrestins and *S. cerevisiae* arrestins*^a^*

*^a^* The blastp and PSI-BLAST programs were used to identify amino acid sequence conservation. Alignments with an E value less than 1 were determined to be significant (N/A = not applicable).

**TABLE S2.** Primary amino acid sequence homology between the *C. neoformans* arrestins and human arrestins*^a^*

*^a^* The blastp and PSI-BLAST programs were used to identify amino acid sequence conservation. Alignments with an E value less than 1 were determined to be significant (N/A = not applicable).

**TABLE S3.** A total of 1,122 proteins were identified as potential interactors of Ali1-GFP*^a^*

*^a^* All identified *C. neoformans* proteins are included, along with the exclusive unique peptide count for each protein in each replicate.

**TABLE S4.** The 59 biologically-relevant proteins identified as potential interactors of Ali1-GFP in YPD medium*^a^*

*^a^* The average peptide count (APC) was calculated by averaging the exclusive unique peptide counts for each potential interactor across three biological replicates. The percent of APC identified in the WT immunoprecipitation was calculated by dividing the APC of each potential interactor by the exclusive unique peptide count found in the WT immunoprecipitation. All potential protein interactors with an APC of 2 or more, as well as less than 20% of the APC identified in the WT immunoprecipitation, were considered to be unique interactors with Ali1-GFP. These potential protein interactors are prioritized by APC (most to least) and percentage of APC identified in the WT control (lowest to highest).

**TABLE S5.** The 62 biologically-relevant proteins identified as potential interactors of Ali1-GFP in TC medium*^a^*

*^a^* The average peptide count (APC) was calculated by averaging the exclusive unique peptide counts for each potential interactor across three biological replicates. The percent of APC identified in the WT immunoprecipitation was calculated by dividing the APC of each potential interactor by the exclusive unique peptide count found in the WT immunoprecipitation. All potential protein interactors with an APC of 2 or more, as well as less than 20% of the APC identified in the WT immunoprecipitation, were considered to be unique interactors with Ali1-GFP. These potential protein interactors are prioritized by APC (most to least) and percentage of APC identified in the WT control (lowest to highest).

**FIGURE S1.** Cellular morphology of the *ali1*Δ mutant at 30°C. The WT, *ali1*Δ mutant, and *ali1*Δ + *ALI1* strains were incubated in YPD medium at 30°C, imaged by DIC microscopy (Zeiss Axio Imager A1), and quantified for the frequency of cytokinesis defects. The percentage of total cells displaying morphological defects at 30°C was quantified for each strain. A minimum of 600 cells were analyzed across three biological replicates (n = 3). Error bars represent the SEM. Log transformation was used to normally distribute the data for statistical analysis (One-way ANOVA; ns = not significant).

**FIGURE S2.** Complementation phenotypes of the individual arrestin mutants. A. Serial dilutions of the WT, *ali2*Δ mutant, and the *ali2*Δ + *ALI2-GFP* strains were incubated on YPD medium, YPD with caffeine (1 mg/mL), and YPD with high salt (1.5 M NaCl). These strains were incubated at 30°C and monitored visually for growth. B. Serial dilutions of the WT, *ali3*Δ mutant, and the *ali3*Δ + *ALI3* strains were incubated on YPD medium incubated at 30°C, YPD medium incubated at 39°C, and YPD with SDS (0.03%) incubated at 30°C. These strains were monitored visually for growth. C. Serial dilutions of the WT, *ali4*Δ mutant, and the *ali4*Δ + *ALI4* strains were incubated on YPD medium and YPD with SDS (0.03%). These strains were incubated at 30°C and monitored visually for growth.

**FIGURE S3.** Virulence contributions of Ali2. The WT, *ali2*Δ mutant, *ali2*Δ + *ALI2-GFP*, and *ali1*Δ*ali2*Δ mutant strains were co-incubated with J774A.1 murine macrophages at a MOI = 1 for 24 hours. Survival of the strains was assessed by quantitative culture, and the percentage of recovered CFU was normalized to the WT strain. This experiment was performed with five biological replicates (n = 5). Error bars represent the SEM. Log transformation was used to normally distribute the data for statistical analysis (*One-way ANOVA p < 0.05; ****One-way ANOVA p < 0.0001; ns = not significant).

**FIGURE S4.** The effects of lipid supplementation on the arrestin null mutants. A. Serial dilutions of the WT and arrestin null mutant strains were incubated on YPD medium at 30°C; YPD at 39°C; YPD with glycerol (0.4%) at 39°C; and YPD with sorbitol (1M) at 39°C. These strains were monitored visually for growth. B. Serial dilutions of the WT and arrestin null mutant strains were incubated on YPD medium; YPD with caffeine (1 mg/mL); YPD with caffeine and glycerol (0.4%); and YPD with caffeine and sorbitol (1M). These strains were incubated at 30°C and monitored visually for growth.

**FILE S1**. Supplementary materials and methods

